# Feasibility of an anterior abdominal electrode array for non-invasive measurement of uterine contractions across the menstrual cycle

**DOI:** 10.64898/2026.10.30.763203

**Authors:** Sophia Shen, Zhenan Bao, Todd Coleman

## Abstract

While traditionally associated with pregnancy and labor, uterine contractions (UC) also occur throughout the menstrual cycle. Abnormalities in UC frequency and/or direction throughout the cycle are correlated with many uterine disorders, which can have downstream impacts on overall health, necessitating continuous monitoring throughout the cycle. Current UC monitoring technologies are invasive or bulky, and not suitable for routine monitoring. The most promising approach uses non-invasive surface abdominal electrodes in a method known as electrohysterography (EHG). However, current EHG methods remain unsuitable for longitudinal monitoring due to bulkiness and reliance on patient-specific anatomical scans. Furthermore, there are no simulations of non-pregnant UC to aid in resolving these limitations. In this work, we evaluate the feasibility of EHG using an electrode array placed only on the anterior abdomen of size 6.35 x 14 cm and without any knowledge of patient anatomy for non-invasive measurement of UC across the menstrual cycle though both simulation and experiment. We identify a signal-to-noise ratio necessary to resolve frequency, amplitude, and direction of UC in simulation. In experimental data, all recordings met the coherence criterion identified in simulation, and we find patterns across the menstrual cycle which are congruent with previous literature. These results demonstrate the feasibility of an anterior abdominal electrode array for monitoring UC across the menstrual cycle, a step towards wearable EHG.

## INTRODUCTION

Although traditionally associated with pregnancy and labor, uterine contractions (UC, also known as uterine peristalsis) also occur throughout the menstrual cycle^1–4^. These contractions arise from the subendometrial myometrium of the uterus, or junctional zone, and vary cyclically across the menstrual cycle in both frequency and propagation direction^3,4^. During menstruation, the contractions are low-frequency (around 1.5 cycles/min) and predominantly in the fundus-to-cervix (F-C), or outward, direction thought to aid in the removal of menstrual blood. On the other hand, during the periovulatory phase, the contractions are high-frequency (around 3 cycles/min) and predominantly in the cervix-to-fundus (C-F), or inward, direction thought to aid in fertilization^2,4^. These coordinated UC play essential roles in reproductive function yet are challenging to measure regularly.

Abnormalities in uterine contraction patterns have been associated with uterine disorders and reproductive outcomes including primary dysmenorrhea^5,6^, secondary dysmenorrhea (e.g., endometriosis)^1,7^, and in-vitro fertilization (IVF)^8,9^. For example, in those with endometriosis, there is an increase in UC frequency and amplitude throughout the entire menstrual cycle, along with a reversal in the direction of the contractions^7,10^. As such, UC have the potential to play an important role in non-invasively assessing reproductive health. Because UC vary throughout the cycle and may differ among individuals, a recording at a single point in time may be unable to identify abnormalities, necessitating longitudinal monitoring. In addition to its clinical utility, continuous monitoring of UC could provide a window into the fundamental physiology of uterine contractile dynamics across the menstrual cycle, offering opportunities for research into mechanisms underlying both normal reproductive function and disease states. Development of an accurate tool for continuous, ambulatory UC monitoring would therefore serve both clinical and basic research goals.

Despite the need for UC monitoring throughout the menstrual cycle, commercially available methods for monitoring UC are tailored toward labor monitoring and lack sensitivity to detect the mild uterine contractions that occur outside of pregnancy. More sensitive methods have been explored for UC monitoring throughout the menstrual cycle, including the intrauterine pressure (IUP) catheter^4,11^ and intrauterine electrodes (IUE)^12,13^. However, these methods are highly invasive and require probe insertion into the uterus. Transvaginal ultrasound (TVUS) is an imaging method that is less invasive, only involving insertion of a probe into the vaginal canal, but results are highly operator-dependent^14,15^. Cine magnetic resonance imaging (MRI) has also been used for measuring uterine contractions throughout the menstrual cycle and has the benefit of being non-invasive; however, it is very expensive, making regular recordings impractical^16,17^. Electrohysterography (EHG) is an established form of monitoring UC in labor and pregnancy^18,19^ and involves electrode arrays placed on the skin around the abdomen. A recent study has shown that EHG can also be used to measure and spatiotemporally categorize UC throughout the menstrual cycle^20^. This method is highly promising, as it is completely non-invasive and relatively inexpensive. Research in the field of non-pregnant EHG is still in its infancy, with only one publication thus far (to our knowledge) demonstrating such accuracy in measuring UC. Barriers to more widespread adoption of current non-pregnant EHG monitoring include: (1) it requires an anatomical MRI scan for source localization, adding substantial cost to the system; (2) it uses 128 electrodes placed around the lower torso and wired to a large data acquisition system, whose bulk restricts recordings to the lab/clinic setting^20^. Altogether, the limitations of current methods highlight their inadequacy for longitudinal monitoring of UC throughout the menstrual cycle, as such devices need to be non-invasive, relatively inexpensive, and compatible for wearable applications (Table 1). Thus, there remains a knowledge gap on whether a wearable-sized EHG electrode array can feasibly measure UC without knowledge of individual anatomy.

**Table 1.** Evaluation of established methods for UC measurement in their ability to measure important UC parameters and satisfy device constraints necessary for continuous measurement. IUP: intrauterine pressure, IUE: intrauterine electrodes, TVUS: transvaginal ultrasound, MRI: magnetic resonance imaging, EHG: electrohysterography.

| **Requirements** | *UC parameter* |  |  | *Device constraint* |  |  |
|---|---|---|---|---|---|---|
| **Measurement Method** | Frequency | Amplitude | Direction | Non-invasive | Relatively inexpensive | Compatible for wearable |
| IUP | Y | Y | N | N | Y | N |
| IUE | Y | Y | Y | N | Y | N |
| TVUS | Y | Y | Y | N | Y | N |
| Cine MRI | Y | N | Y | Y | N | N |
| Previous EHG | Y | Y | Y | Y | N | N |

When there exist many unknowns in a system which are challenging to directly control or observe, simulation is a method to predict expected trends. As compared to in vivo recordings, simulation of UC enables independent variation of factors such as noise level, anatomy, and wave type to compare resulting measurements against the ground truth. Existing computational models of UC focus on labor and pregnancy, where the signals have different spatiotemporal characteristics from those during menstrual cycle. While these models incorporate aspects such as electrophysiology and volume conduction through surrounding tissues^21–23^, they do not account for the unique anatomy and physiology pertaining to the non-pregnant state. Thus, there remains another knowledge gap, motivating the development of a simulation framework specifically for EHG signals of UC throughout the menstrual cycle.

Here, we address both knowledge gaps. First, we do this by presenting what is, to our knowledge, the first anatomically based volume-conduction model of UC outside of pregnancy. We use the simulation to evaluate feasibility of using a small electrode array (6.35 x 14 cm) on the anterior abdomen for recovering spatiotemporal characteristics at various noise levels. Then, we assemble a custom electrode array of the same size and measure UC in healthy participants across all four phases of the menstrual cycle. In human subjects, we next assess the feasibility of extracting amplitude and spatiotemporal information about the contractions across the menstrual cycle. Altogether, this study evaluates the feasibility of using an anterior abdominal electrode array to monitor UC across the menstrual cycle without any knowledge of individual anatomy, a step toward the creation of wearable EHG.

## ANALYSIS METHODS

Both experimental and synthetic data used the same analysis pipeline. Values were averaged across recordings within each phase or day. Significance testing was accomplished using RM-ANOVA with Holm-corrected paired t-tests.

### Frequency and Amplitude values

The dominant frequency for each recording was determined by calculating the mean peak frequency from a Welch power spectral density (PSD) across electrodes. The amplitude was determined using root mean square (RMS), which was computed over the full recording for each electrode, then averaged across electrodes. These methods are similar to previous publications of electrogastrography (EGG), surface electrophysiology measuring contractions of the stomach, where the waveforms have similar frequencies and wave types^24–26^.

### Direction measures

The overall direction was measured using the phase gradient directionality (PGD) of the signal. This is an established method used to identify direction of wave propagation in EGG^27^, where the direction at each point in time can be calculated using:

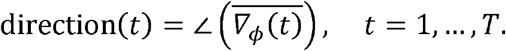

where *∇*_*ϕ*_ is the spatial gradient of the instantaneous phase (computed via the Hilbert transform). How aligned the array is in the identified direction of the wave can then be identified using the PGD value, which is given by:

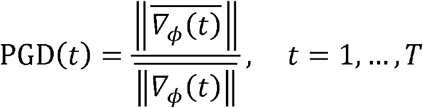

The PGD value ranges from 0 to 1, with 0 being no aligned wave and 1 being a perfect plane wave.

For each recording, the PGD value and direction were identified at every point in time. To determine the most optimal PGD value, we analyzed the angular dispersion of the identified direction values. Using simulated data at a range of signal-to-noise ratio (SNR) values, variation of more than 45° in direction was set as the threshold for when the wave was no longer aligned. As shown in Figure 2a, at SNRs where mean angular dispersion stayed below 45° (≥ −35 dB), 95% of PGD values exceeded 0.113 for both inward and outward. Thus, a wave was considered coherent if the PGD was over 0.15 for 2 seconds, with 0.15 used as a safety margin^27^.

For each recording, the dominant axis (90°/270° or 0°/180°) was identified once for the entire recording by visual inspection, blind to menstrual phase. In each coherent time point, the direction was assigned inward or outward depending upon the menstrual cycle phase and which ratio was larger. This is because in simulated data, we found that which identified direction was actually inward or outward differs based on individual anatomy (Fig. 2b,c). All other non-coherent waves were assigned as “other”.

## RESULTS

### Assessing feasibility of an anterior abdominal electrode array for EHG through simulation

We first evaluated the feasibility of using an anterior abdominal electrode array for EHG using a finite-element model (FEM) simulation of EHG (Fig 1a-b), adding noise to synthetic datasets to achieve a range of SNR values (Fig 1d).

**Figure 1.**
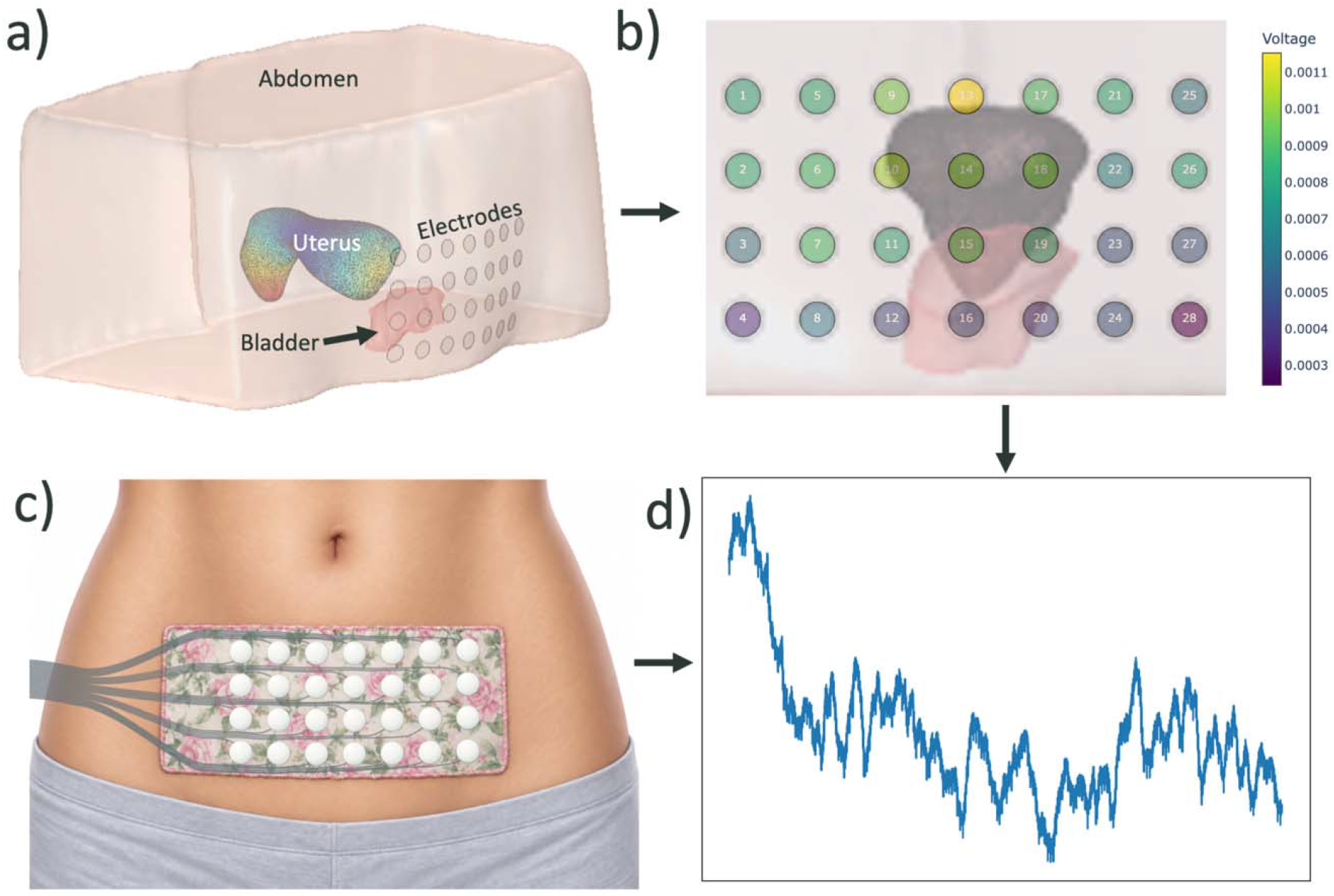
Pipeline for simulated and experimental data. Simulated pipeline: a) 3D models of the uterus, abdomen, and bladder were imported into the FEM model, where a dynamic wave was simulated along the medial curve of the uterus. b) Solution of forward model incorporating volume conduction using FEM to generate the dynamic simulated EHG signal. Experimental pipeline: c) Participants wore a custom electrode array on the anterior abdominal surface. d) Exemplar waveform for both methods with measurement noise, from one electrode in the array (AWGN added to simulated data from (b), and raw data acquired from (c)).

**Figure 2.**
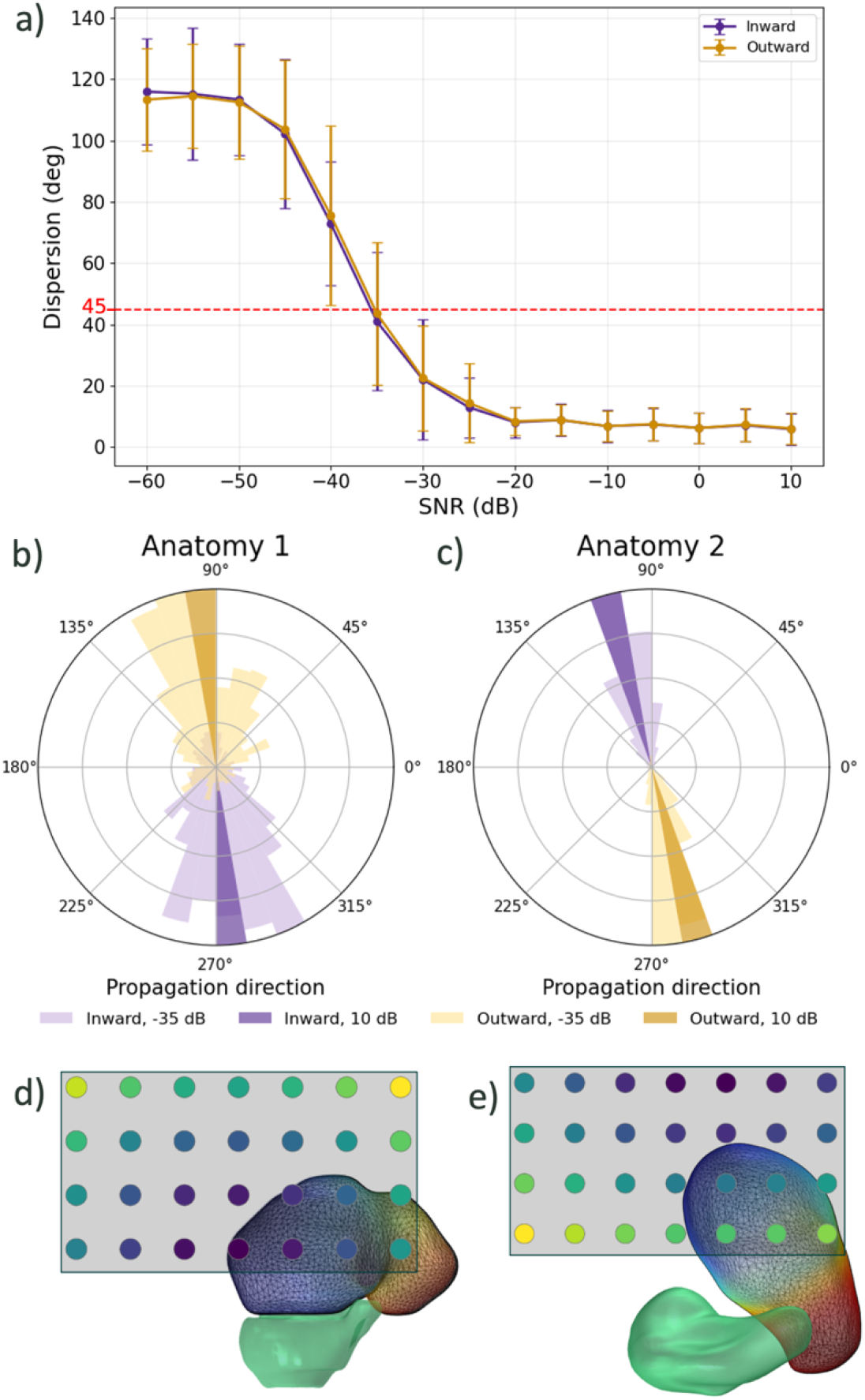
Determination of methods for direction analysis. a) The mean angular dispersion of the classified direction of simulated inward and outward waves at various SNR levels, with SD error. The red dashed line marks 45° of dispersion, below which is acceptable variation in the direction. b,c) Spatial histograms visualizing the angular dispersion in 2 sample anatomies at lowest optimal (-35 dB) and a very good (10 dB) SNR. d,e) 3D model demonstrating why different anatomies resulted in opposite assigned propagation directions. Darker colors represent lower voltage (uterine surface) or phase (electrode array).

First, we examined how the identifiability of frequency changed with SNR (Fig. 3a). With very low SNR values, the mean identified frequency was much higher than the actual, with a large variation of 0.68 cycles/min. Starting at -35 dB for all four phases of the menstrual cycle, the identified frequency closely matched the actual frequency, with much lower standard deviation of 0.05 cycles/min. Thus, an SNR of at least -35 dB is necessary for identifying the correct frequency.

**Figure 3.**
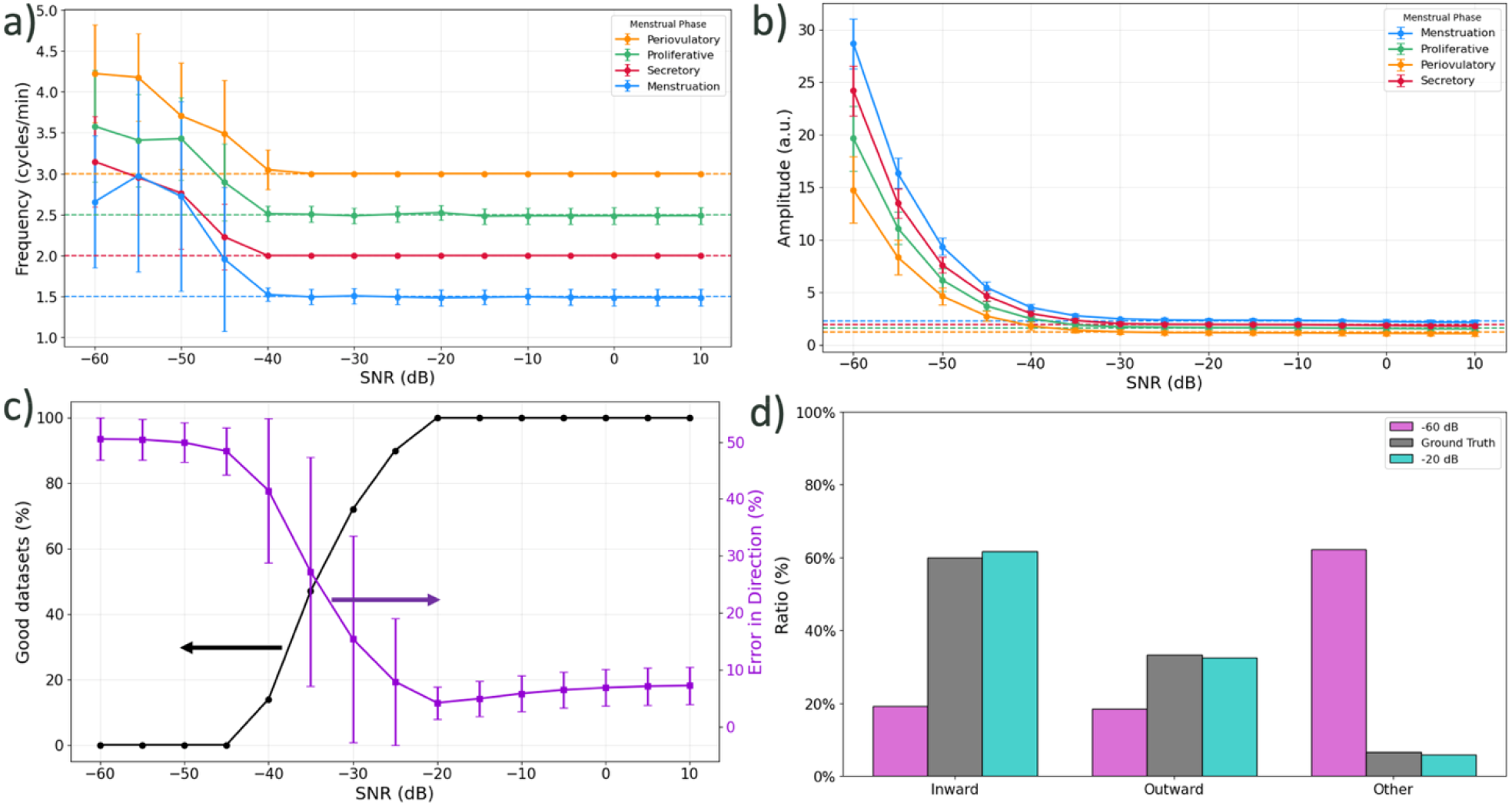
Simulated dataset results across SNR. a) Identified frequency of datasets at various SNR levels in all four menstrual cycle phases, with dashed lines representing the true frequency value of the dataset. b) Amplitude of datasets at various SNR levels in all four menstrual cycle phases. c) The percentage of physiologically feasible, good datasets (black) and the mean error in assigned direction ratios (purple) at various SNR levels. d) The assigned ratio of inward, outward, and other waves in a sample synthetic dataset from the periovulatory phase at very bad (-60 dB, pink) and acceptable (-20 dB, turquoise) SNR levels. All error bars represent standard deviation.

We next measured the effects of SNR on amplitude (Fig. 3b). Similar to frequency, the magnitude of error in amplitude measurements decayed monotonically with SNR. As there is no consensus on the absolute amplitude of uterine contractions^28,29^, the values are presented in arbitrary units. The relative error at these low SNR values was inversely related to the frequency values, with the lowest frequency (menstruation) having the highest error of 26, while all four phases had similar standard deviations of 2.73. In all four phases of the menstrual cycle, the difference between identified amplitude and actual amplitude was the lowest at SNR values of at least -25 dB, reflecting a minimum SNR of -25 dB to correctly evaluate amplitude.

Finally, we explored the ability to identify direction correctly with SNR. We began by determining how SNR affected the percentage of datasets were physiologically feasible (“good”), which was defined according to previous literature as having more than 65% of the waves being coherent (inward or outward)^11,20^. Since these synthetic datasets were all generated containing more than 65% of coherent waves, the expected percentage of good datasets was determined to be 100%. Similar to amplitude, we also measured the effects of SNR on the error in assigned direction, comparing the average percent error in predicted direction proportions. As shown in Figure 3c, both the percentage of good datasets reached 100% and the error plateaued at -20 dB. This demonstrates a minimum SNR for reliable retrieval of direction. Figure 3d shows the assigned direction proportions for a sample synthetic dataset for the periovulatory phase, demonstrating that in high noise situations, many coherent (inward or outward) waves are assigned as non-coherent (other) waves, leading to increased error.

### Assessing feasibility of an anterior abdominal electrode array for EHG through experimental recordings

We recorded EHG from 20 participants, once in each phase of the menstrual cycle (80 recordings), using a custom 28-electrode array measuring 6.35 × 14 cm (Fig. 1a). All 80 recordings met the criterion for physiologically feasible datasets (more than 65% coherent waves).

First, we plotted the frequency of the EHG across the menstrual cycle, both by day (Fig. 4a) and by menstrual cycle phase (Fig. 4b). Scaled across a standard 28-day cycle, the highest frequency occurred mid-cycle around days 15-17, with the lowest frequency early in the cycle. By menstrual cycle phase, there was a significant difference in frequency across phases, with the highest corresponding to the periovulatory phase and the lowest corresponding to the menstruation phase. A similar but non-significant pattern was seen by phase for amplitude, where the highest occurred during the periovulatory phase and the lowest occurred during the menstruation phase (Fig. 4c). Analysis of the ratio of the wave directions across the menstrual cycle by day showed the expected ratio of inward and outward were inversely related, with outward being more predominant during the first half of the cycle and inward during the second half (Fig. 4d). The two curves crossed over just before mid-cycle. Furthermore, the ratio of other waves did not change much in comparison.

**Figure 4.**
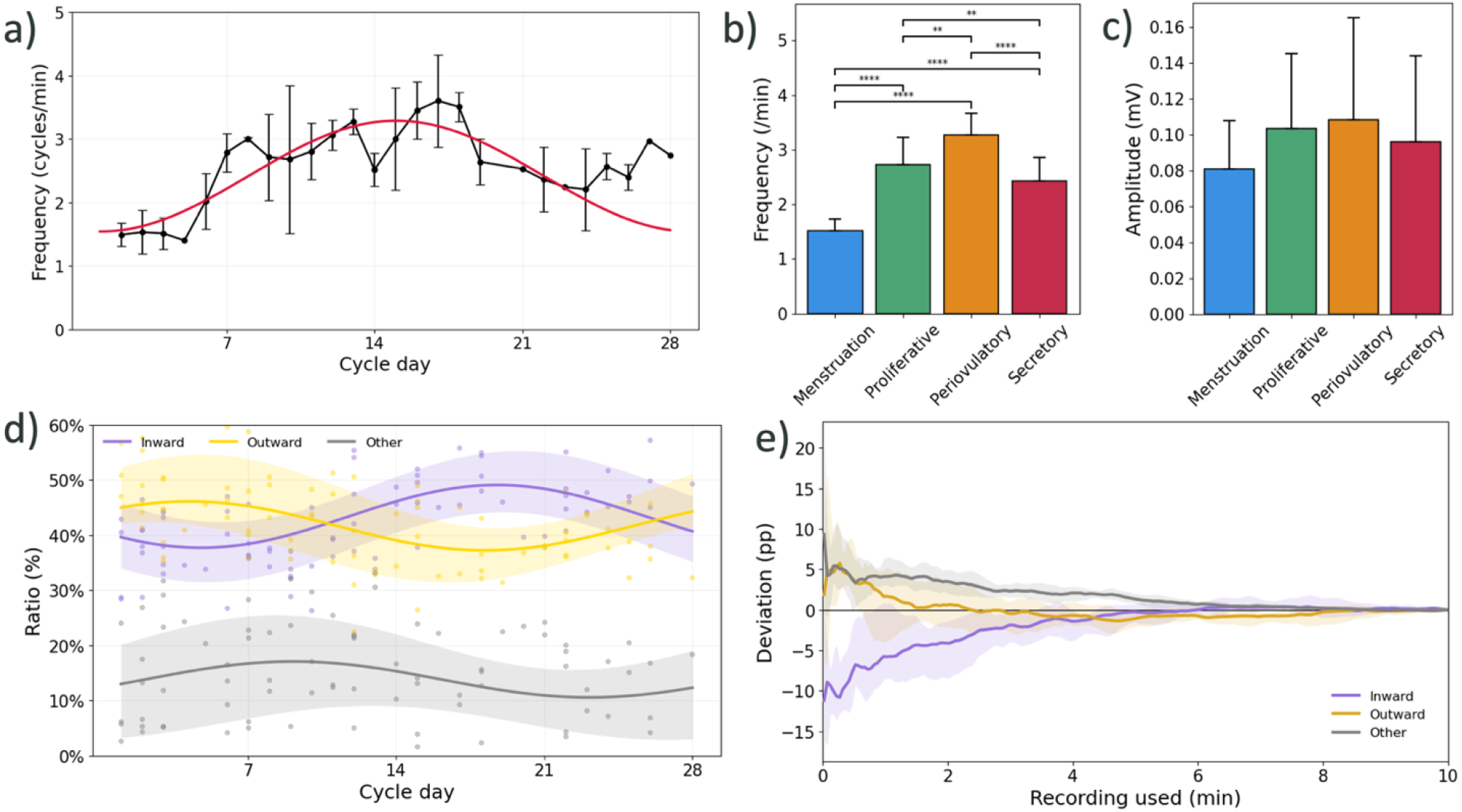
Experimental results across the menstrual cycle. a) Frequency of UC by day normalized across a 28-day cycle. Red line represents a regression fit with Huber robust least-squares using the means. Significance is marked as ** p < 0.01, **** p < 0.0001. b) Frequency of UC by menstrual cycle phase. c) Amplitude of UC across each phase of the menstrual cycle. d) Assigned ratio of inward, outward, and other waves for each recording across the menstrual cycle by day, normalized across the 28-day cycle. Mean curves were fit using harmonic regression with robust loss described in (a); shaded bands were fit using the same method applied to the daily mean ± SD. Dots represent individual values e) Deviation in percentage points of the calculated cumulative direction ratio from the direction ratio calculated using the entire recording for each of the directions (inward, outward, and other). Shaded bands represent SD, x-axis scaled to a nominal 10-minute recording.

To better characterize the relationships between wave directions across the menstrual cycle, a first-order Markov chain was modeled for each recording by tallying transitions between the states of consecutive windows, which was then normalized by total counts to give a transition probability for each state (Table 2). Among transitions between states, those toward the phase-dominant direction were significantly more probable: outward in the first two phases and inward in the last two. Additionally, self-transitions had the highest transition probabilities, indicating that direction persisted across consecutive windows.

**Table 2.** Transition probabilities for Markov chain model of wave directions (inward, outward, other) over time. Omnibus p-values are from RM-ANOVA across the four phases. Significance is marked as * p < 0.05, ** p < 0.01.

| Transition |  | Omnibus | Transition probability |  |  |  |
|---|---|---|---|---|---|---|
| From | To | p-value | Menstruation | Proliferative | Periovulatory | Secretory |
| Inward | Inward | * 0.0114 | 86.3% | 84.8% | 88.3% | 87.7% |
| Inward | Outward | ** 0.0089 | 8.2% | 9.1% | 7.0% | 7.5% |
| Inward | Other | 0.3994 | 5.6% | 6.1% | 4.7% | 4.9% |
| Outward | Inward | ** 0.0075 | 6.7% | 7.0% | 8.7% | 8.3% |
| Outward | Outward | * 0.0106 | 88.3% | 87.1% | 84.7% | 85.6% |
| Outward | Other | 0.3132 | 4.9% | 5.9% | 6.6% | 6.1% |
| Other | Inward | ** 0.0061 | 13.2% | 12.5% | 17.0% | 16.4% |
| Other | Outward | 0.0943 | 17.9% | 15.7% | 14.1% | 15.0% |
| Other | Other | 0.1598 | 68.8% | 71.8% | 68.9% | 68.6% |

Finally, we aimed to assess the minimum length of recording needed for successful extraction of spatial parameters. This was done by assessing direction ratios only using cumulative portions of each recording and comparing it to the final ratio, starting from 2 seconds to the full recording. As shown in Figure 4e, with the shortest recording there was around 5 pp deviation on average for outward and other waves, and around -10 pp for inward waves. As the recording data became longer, the error decreased, stabilizing at around 8 minutes for all directions with less than 0.25% error.

## DISCUSSION

Overall, the results presented in this paper demonstrate feasibility of using an anterior abdomen electrode array to measure frequency, amplitude, and direction throughout the menstrual cycle without subject-specific anatomical knowledge, which, to our knowledge, has not been done previously.

The results from the simulated data demonstrate a minimum SNR for measuring spatiotemporal characteristics of UC of -20 dB. Frequency analysis was the most robust to noise, having a low threshold of -35 dB, due to the fact that much of the noise occurs at higher frequency bands, which is removed by our filtering process. The error in amplitude was also found to be inversely related to the frequency of the signal at low SNR. This may reflect the nature of the Wiener filter (as described in Methods section), where lower frequencies translate to higher window lengths and consequently fewer windows per recording to average over, which increases the error. However, in settings with SNR above the identified threshold of -25 dB, this effect disappears. With direction metrics, -20 dB was the identified threshold for retrieving direction metrics with no datasets removed for poor data quality and the lowest error. Thus, we deem -20 dB the lowest SNR for which UC characteristics are reliably retrievable within the context of this in silico paradigm. To assess whether our experimental recordings meet this criterion, we applied the same PGD-based coherence threshold (Fig. 3c) and found that 100% of recordings satisfied it, the feasibility standard established via simulation.

Using the methods validated from the simulated results, analysis of the spatiotemporal characteristics in the experimental data yielded results consistent with previous literature. Specifically, the changes in frequency throughout the cycle were significant, with the lowest during menstruation and the highest during periovulatory. Although there were variations in the frequency per day, these may reflect our limited dataset and natural fluctuations throughout the cycle during the day, which remains poorly studied^30,31^. In terms of amplitude, we found a similar pattern to frequency with menstruation being the lowest and periovulatory being the highest, which is consistent with what some previous studies had also observed^32,33^. However, it is worthy to note that the amplitude of contractions is not widely agreed upon, with different studies noting different patterns, and the difference may be due to the measurement method^11,32– 35^. In terms of direction, coherent propagation was identified in all recordings, with non-coherent waves making up only a minority of windows, and waves tended to remain in the same direction across consecutive windows. Using a phase-based labeling rule necessitated by variations in individual anatomy, inward and outward proportions followed the pattern reported in previous literature^10,14^, with outward waves more frequent early in the cycle and inward waves later.

The length of recordings for the experimental study for each session was approximately 10 minutes. In the past, other studies have had varying times, ranging from 2 to 30 minutes^35,36^. Thus, there is no standardization on the optimal recording length, which may also differ between UC recording methods. We found that 8 minutes was sufficient for our recording on average, as the direction ratio was stable for the remainder of the recording. This suggests that 10 minutes was sufficient for this array and analysis and that approximately 8 minutes may be enough to estimate direction ratios.

This paper demonstrates the feasibility of using wearable-size electrode arrays for EHG measurement of frequency, amplitude, and propagation direction. These arrays have several advantages over previous methods of monitoring UC. First, no invasive exams or expensive subject-specific anatomical scans are necessary – the entire system functions using measurements from the skin surface. Second, the array’s small footprint is comparable to existing wearable electrode systems^37^, making a wearable EHG device a plausible next step. Third, the array is able to obtain amplitude and spatiotemporal information about the contractions, which not all previous methods could offer. Together, these advantages provide the potential for an ambulatory EHG system to enable continuous monitoring over the menstrual cycle, which could provide valuable insights into day-to-day and cycle-to-cycle changes important for reproductive health.

There were some limitations of this study that are worth mentioning. First, the simulation is a simplification of the real world, which contains physiological signals from other sources such as the heart, bladder, and stomach. However, these signals should have an attenuated effect on the results, as cardiac activity is ∼10x higher in frequency and the bladder was empty or near-empty. For the stomach, the time of the last meal was available for 67 of 80 recordings, and amplitude in the gastric frequency band (0.03–0.07 Hz) was not significantly different between recordings made within 1, 2, or 3 hours of a meal and later recordings. Given that gastric amplitude typically increases for multiple hours after meal consumption^24,38,39^, this suggests that our recordings are not of gastric origin. Second, the inward or outward direction of propagation in the experimental recordings could not be verified against a ground truth. Because the identified angle alone does not specify absolute direction, inward and outward labels were assigned using a phase-based rule. Third, both the simulated and experimental datasets have limited number of individuals, which limits generalization but provides a foundation for feasibility.

## FUTURE DIRECTIONS

Future work will build upon these results. On the simulation side, we hope to extend simulation capabilities and amass a larger dataset of 3D models, which would allow us to develop a machine learning model to classify specific wave types and help determine propagation directions from a single recording. On the experimental side, we plan to develop the array into a wearable device that can enable continuous monitoring outside of the lab setting, along with validation using simultaneous recording. This would allow for further large-scale studies that can examine many unanswered questions in this field, including how UC change throughout the day, as well as clinical studies examining correlations of UC with uterine disorders. Such a device could establish UC as a biomarker of reproductive health.

## METHODS

### Finite-element simulation of EHG

MRI data (axial, sagittal, and coronal views) were obtained from the publicly available SynthRAD dataset^40^. Segmentation of the bladder, uterus, and abdomen from these datasets was performed using 3D Slicer^41,42^ (Fig. 1a). The segmented 3D models were then imported into COMSOL for simulation. The abdomen was then replaced with an elliptical cylinder matching the major and minor axes of the abdomen to reduce complexity, as it has been found that increasing complexity of the model does not result in a large increase in accuracy^43^. The electrical potential along the uterus along the medial curve was simulated using the 1D wave equation^44,45^. The resulting waveforms were then propagated to surface abdominal electrodes using volume conduction, with electrical conductivity of the abdomen as 0.125 S/m (homogenized between abdominal muscle, adipose tissue, and skin), uterus as 0.464 S/m, and bladder as 0.5 S/m to represent an empty bladder (Fig. 1c). The simulated electrode array matches the size and shape of the experimental EHG electrode array (see below).

### Generation of synthetic dataset

Synthetic datasets for each phase of the menstrual cycle were created by stitching together single-cycle segments extracted from simulated template recordings to form 10-minute recordings. Each template recording corresponded to one of five propagation types — inward, outward, converging, diverging, or standing — generated at one of four frequencies corresponding to menstrual cycle phase (menstruation: 1.5 cycles/min; proliferative: 2.5 cycles/min; periovulatory: 3 cycles/min; secretory: 2 cycles/min). The relative ratios of the propagation types were randomly drawn from ranges corresponding to previously reported ratios of inward, outward, and other (converging, diverging, standing, or pure noise created separately from simulation) waves occurring in each phase of the menstrual cycle^20,46^. Five synthetic datasets were created for each anatomy/phase combination, for a total of 100 different synthetic datasets. Broadband additive white Gaussian noise (AWGN) was added to each dataset to achieve target signal-to-noise ratios.

### Participant enrollment

This study was approved by the Stanford University Institutional Review Board, and all participants signed informed consent documents. Participants were included if they were between the ages of 18-40 and had a regular menstrual cycle (defined as between 21-35 days). Potential participants were excluded if they were pregnant, used any form of hormonal birth control, had been diagnosed with any uterine abnormalities, or had any self-reported histories of menstrual cycle abnormalities. The experimental data from this study were obtained from 20 healthy non-pregnant pre-menopausal participants. The detailed demographics of the participants can be found in Table 3. The study was approved by the Stanford IRB, and all participants provided written informed consent.

**Table 3.** Demographics from experimental EHG study (mean ± SD).

| **Age (years)** | 25.85 ± 3.47 |
|---|---|
| **BMI (kg/m<sup>2</sup>)** | 21.87 ± 2.38 |
| **Menstrual Cycle Length (days)** | 29.05 ± 3.15 |
| **Race (n, %)** |  |
| Asian | 13, 65% |
| White | 4, 20% |
| Black | 2, 10% |
| Mixed | 1, 5% |

### Experimental EHG data collection

EHG data was collected using a custom-made 4x7 electrode array consisting of 28 BrainVision actiCAP slim electrodes. Electrodes were spaced 1.5 cm apart. The array was placed on the lower abdominal surface, with its lower edge above the pubic hairline, centered on the midline, as the anterior lower region has been shown to provide the most information^47^. Data was recorded using a BrainVision actiCHamp amplifier, sampled at 1000 samples/second, with each recording being 10 minutes. Participants were asked to empty their bladder prior to each recording. For each participant, a recording was taken for each phase of the menstrual cycle, for a total of four recordings per participant. Menstrual phase was determined based on self-reporting of menstruation and urine-based luteinizing hormone (LH) ovulation tests.

### Electrophysiological signal preprocessing

Both experimental and synthetic data used the same signal preprocessing pipeline (Fig. 1d), with the exception that common average referencing (CAR) was applied to remove widespread, correlated noise across all electrodes in experimental data only. Next, 8^th^ order Butterworth notch filter was used to remove 60Hz line noise. Then, a Wiener filter was used for adaptive artifact rejection (a method previously utilized and optimized for EGG^24^), where the window size input varied based on reported frequency values for each phase of the cycle. Finally, a 4^th^ order Butterworth bandpass filter from 0.01 to 0.1 Hz was applied to filter the final signal to the frequency range of interest, and transients were removed via trimming.

## AUTHOR CONTRIBUTIONS

S.S. designed the experiment, collected and analyzed data, and wrote the manuscript. All authors reviewed and edited the manuscript.

## ACKNOWLEDGEMENTS

This work was supported in part by Stanford University’s Community of Shared Research Platforms (C-ShaRP), the Stanford Wearable Electronics Initiative (eWEAR) seed funding, and Tianqiao and Chrissy Chen Ideation and Prototyping Lab. We gratefully acknowledge the use of facilities, equipment, and voucher funding provided by the program. Author S.S. was supported by a National Science Foundation Graduate Research Fellowship, National Institutes of Health Predoctoral Individual National Research Service Award (F31), Stanford BIO-X SIGF Fellowship, and the Wu Tsai Human Performance Alliance Honorary Fellowship at Stanford University. Author T.P.C. was supported by the Wu Tsai Human Performance Alliance. We would like to thank Katelyn Yeh for assistance with segmentation and simulation. We would also like to thank Suchita Rao and Sarah Arnold for assistance with signal processing.

## DATA AVAILABILITY

The data reported in this work are available from the corresponding author upon reasonable request.

## CODE AVAILABILITY

The underlying code for this study are available from the corresponding author upon reasonable request.

